# Fingerprinting CANDO: Increased Accuracy with Structure and Ligand Based Shotgun Drug Repurposing

**DOI:** 10.1101/591123

**Authors:** James Schuler, Ram Samudrala

## Abstract

We have upgraded our Computational Analysis of Novel Drug Opportunities (CANDO) platform for shotgun drug repurposing to include ligand-based, data fusion, and decision tree pipelines. The first version of CANDO implemented a structure-based pipeline that modeled interactions between compounds and proteins on a large scale, generating compound-proteome interaction signatures used to infer similarity of drug behavior; the new pipelines accomplish this by incorporating molecular fingerprints and the Tanimoto coefficient. We obtain improved benchmarking performance with the new pipelines across all three evaluation metrics used: average indication accuracy, pairwise accuracy, and coverage. The best performing pipeline achieves an average indication accuracy of 19.0% at the top10 cutoff, compared to 11.7% for v1, and 2.2% for a random control. Our results demonstrate that the CANDO drug recovery accuracy is substantially improved by integrating multiple pipelines, thereby enhancing our ability to generate putative therapeutic repurposing candidates, and increasing drug discovery efficiency.

## Introduction

### Drug repurposing

Bringing a new drug to the market may costs hundreds of millions of dollars and takes years of work.^1^ Drug repurposing is the process of discovering a new use for an existing drug.^2,3^ This process may take advantage of existing data on safety and pharmacokinetic properties from previous trials and clinical use to reduce costs and time associated with traditional drug discovery. Classic examples of drug repurposing include sildenafil (Viagra) and thalidomide, which initially were developed to treat chest pain and morning sickness, but were repurposed to treat erectile dysfunction and erythema nodosum leprosum respectively.^2,4,5^ Drugs which have already been repurposed once are being researched for even more novel uses. For example, raloxifene was originally indicated for prevention of osteoporosis and was subsequently approved for risk reduction in the development of breast cancer. ^6^ More recently, raloxifene has been suggested as a possible treatment for Ebola virus disease^7–9^. These examples of putative and/or successful drug repurposing underlies the diverse mechanisms through which a single compound may treat a variety of disease types.^10,11^ High throughput, target-based, and phenotypic screening of compounds can be used to generate putative candidates for repurposing.^12^ For example, potential treatments for Zika virus infection were identified using a phenotypic screen.^13^

### Computational drug discovery and repurposing

Finding new drugs or new uses for existing drugs computationally takes advantage of the growing amount of data generated from wet lab experiments accessible on the Internet, increased computational power, and higher fidelity of computational models to reality. Approaches to computational drug discovery and repurposing have been classified as structurebased or ligand-based.^14–16^ In structure-based methods, the structure of a target macro-molecule, usually a protein, is used to identify small compounds that modulate its behavior. The structure may have been determined via x-ray diffraction or Nuclear Magnetic Resonance (NMR), or modeled using template-free (*de novo*) or template-based (homology) modeling.^17–19^ Molecular docking and/or rational drug design is then used to identify ligands that specifically fit into a protein groove or active site. ^20,21^ In ligand-based methods, the focus is on the compound, and similarity between representations is used to assess whether a compound modulates the activity of a target or treat a disease like a known drug. Examples of ligand-based drug design include 2D and 3D similarity searching,^22^ pharmacophore modeling,^23^ and quantitative structure activity relationships (QSAR).^14^

Data fusion is a technique in the field of cheminformatics for combining intermolecular similarity data from different sources or methods. ^24–26^ Compounds are ranked relative to each other based on the similarity scores. Multiple rankings of compounds produced by different methods of detecting similarity may be combined into a single ranking.^24^ Ideally, disparate sources or types of data may yield orthogonality or complementarity in results, i.e., different top compounds are captured and reported as putative therapeutics for different reasons.^27,28^ For example, Tan et al. obtained an increased recall rate in a virtual screening experiment using ligand-based two dimensional fingerprint data fused with structure-based molecular docking energies.^29^ Ligand-and structure-based methods have been combined for use in virtual screening pipelines and platforms, with successes reported in the use of sequential, parallel, and hybrid techniques for data integration.^28^ Data fusion has been also been used to devise weighting schemes for correct dosing.^30^

Newer computational techniques for drug discovery and repurposing gaining in prominence go beyond the structure and ligand-based categorization. The Connectivity Map is a “reference collection of gene-expression profiles from cultured human cells treated with bioactive small molecules”,^31^ i.e., a tool to identify changes in gene expression due to a compound or a disease. If a compound causes changes in gene expression level opposite to a disease (for instance, a disease causes up-regulation of the expression of a set of genes, and the compound causes down-regulation of the same set of genes), then that compound is considered to be therapeutically useful in the treatment of that disease.^31^ Peyvandipour et al. combined an updated version of the Connectivity Map with knowledge of drug-disease gene networks, measuring the perturbation effect of drugs on whole systems. Using this model, they predicted novel treatments for idiopathic pulmonary fibrosis, non-small cell lung cancer, prostate cancer, and breast cancer, while simultaneously improving the recall rate of known drug-disease associations.^32^ Machine learning based approaches have also been used to cluster drugs or diseases and predicting new drug activity and usage.^33–37^Methods for finding novel uses of drugs based on analysis of biomedical literature,^38,39^ electronic health records,^37,40^ and biological networks^41,42^ have also been reported.

### Drug similarity

Implementations of drug discovery and drug repurposing sometimes rely on the principle of similar molecules having similar properties. ^43,44^ In drug design, repurposing, or screening, similar compounds are generally assumed to have similar molecular targets. In structure-based drug discovery, if two potential molecular targets are identified as similar, then a compound that modulates one target is inferred to modulate the other. In ligand-based methods, similar compounds are inferred to analogously modulate the behavior of the same target(s). In our computational shotgun drug repurposing experiments, we extend the similarity property principle to examining interactions on a proteomic scale. Compounds with similar proteomic interaction signatures are hypothesized to be effective for the same indication(s).

### Shotgun drug repurposing with CANDO

The goal of the Computational Analysis of Novel Drug Opportunities (CANDO) platform for shotgun drug discovery and repurposing is to screen every human use compound/drug against every indication/disease.^45–48^ The tenets of CANDO include docking with dynamics, multitargeting, and shotgun repurposing, which have been developed over the last decade and a half.^49–51^ The first version of CANDO (v1) applied a bioinformatic docking protocol on large libraries of compound and protein structures. The multitargeting nature of drugs^52^ is captured by inferring their similarity on a proteomic scale after calculating interactions between all compounds and all proteins in the corresponding libraries. ^8,45,46^ This is key, as indications can be multifactorial in nature, involving disparate or intertwined pathways.^53**?** –56^ Similar compounds, as determined by the root mean square deviation (RMSD) of their proteomic interaction signatures, are hypothesized to behave similarly, i.e., compounds which are ranked highly (most similar compound-proteome interaction signatures) to a drug with an approved indication are hypothesized to be repurposable drugs/compounds for that indication. Benchmarking is accomplished by examining the ranks of other approved drugs for the same indication.^45,46^

There exist other approaches to determine compound similarity without the need for docking calculations. Different representations of molecules capture different chemical, physical, or functional aspects of a compound. Two or three dimensional molecular fingerprints are used in the field of cheminformatics to describe compounds.^57^ In these models, the physical arrangement of atoms in a compound is captured as a binary vector where each entry of a vector indicates the presence or absence of a specific molecular feature.^44^ A distance (similarity) metric between these vectors can be measured, using metrics such as the Tanimoto coefficient, a widely used metric in medicinal chemistry and ligand-based virtual screening.^44,58–60^ This is analogous to the structure-based methods used to construct interaction signatures in v1 and the RMSD measure used to calculate similarity.

In this study, we extend CANDO to include ligand-based drug repurposing by creating new pipelines based on identifying compound similarity based on their molecular fingerprints, as well as data fusion pipelines that combine the protein-centric and protein-agnostic approaches. The new ligand-based pipelines in CANDO are based on molecular fingerprint similarity calculations using the Research Development Kit (RDKit), ^61^ and are not meant as an exhaustive exploration of all possible CANDO pipelines that can be built using all the fingerprint descriptions available from RDKit. Instead, we constructed pipelines using well studied molecular fingerprints^62^ to evaluate feasibility and compare and contrast benchmarking performance. Using the standard CANDO benchmarking procedure (see “Methods”), several of the pipelines described here yielded better performance than those previously obtained using v1 by itself.

Combination of other pipelines using data fusion as well as a decision tree approach between v1 and the best performing ligand-based approach (“ECFP4”) yielded better benchmarking performance than using either pipeline by itself, allowing for increased accuracy while retaining the mechanistic and precision medicine opportunities afforded by the protein-centric approach of v1. Higher benchmarking accuracies are indicative of higher drug repurposing potential, increased confidence in our predictions, a decreased number of compounds which must be tested in wet lab experiments and clinical trials to obtain true hits, and thus less time and cost required to find a new use for an old drug.

## Methods

Figure 1 illustrates the different pipelines evaluated in this study, which are described in detail below.

**Figure 1:**
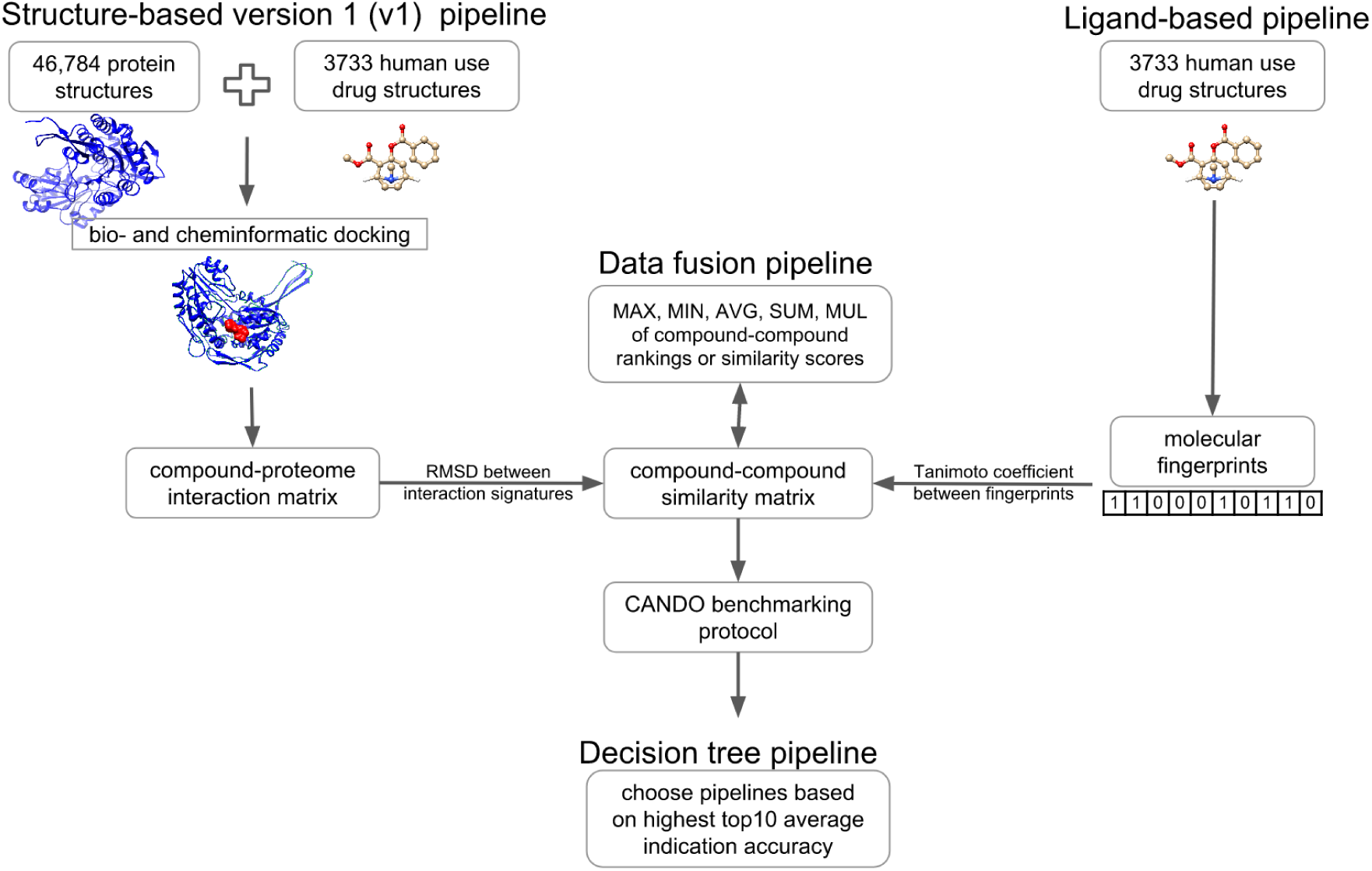
Flow diagram of the CANDO platform pipelines used for shotgun drug repurposing. The v1 structure-based pipeline is the original protein-centric approach based on a bioinformatic docking protocol used to construct compound-proteome interaction signatures. The ligand-based pipelines are based on molecular fingerprint representations of compounds. The data fusion pipelines consist of a combination these two types of pipelines after calculating compound-compound similarity, and the decision tree pipeline is devised based on the performance of individual structure- and ligand-based pipelines (see Methods). All pipelines, except the decision tree pipeline, generate a compound-compound similarity matrix that is sorted and ranked. These rankings are used to generate putative repurposable drug candidates and evaluate benchmarking performance. The figure illustrates the utility of implementing, as well as comparing and contrasting, multiple (types of) pipelines in the CANDO platform for shotgun drug repurposing.

### The CANDO platform and the version 1 (v1) pipeline

A detailed description of the CANDO platform, including the v1 pipeline used for assigning drugs to indications, as well as its benchmarking performance, is available elsewhere. ^45–47,63^ Briefly, in v1 we predicted interactions between 46784 protein structures and 3733 small molecules that mapped to 2030 indications. We obtained the molecular structures of the 3733 small molecules in our putative drug library from the Food and Drug Administration (FDA), NCATS Chemical Genomics Center, and PubChem. ^64^ Solved x-ray diffraction structures of proteins were obtained from the Protein Data Bank ^65^ and modeled protein structures were generated using I-TASSER. ^19^ Approved drug-indication associations were obtained from the Comparative Toxicogenomics Database (CTD)^66^ and mapped to the CANDO drug library, resulting in 2030 indications with at least one approved/associated compound. Protein-compound interaction scores were calculated using a bio- and chem-informatic docking protocol consisting of ligand binding site identification for all proteins in our structure library, followed by similarity measurement between known ligands in the identified binding sites and all 3733 compounds in our putative drug library. ^46^ A compound is characterized as an “interaction signature” of length 46784, where each entry is an interaction score between 0 - 2, indicating the strength of a predicted protein interaction (zero signifying no interaction). Each compound is then compared to every other compound by calculating the root mean square deviation (RMSD) between the corresponding interaction signatures, generating a compound-compound (or drug-compound) similarity matrix. Each compound is ranked relative to every other compound in order of increasing similarity and benchmarking performed.

### Ligand-based pipelines

The CANDO platform for shotgun drug repurposing is not dependent on any particular method for determining compound similarity, such as the protein-centric one used in v1. Here, we consider the utility of ligand-based pipelines by constructing two dimensional molecular fingerprints of the 3733 compounds in the CANDO putative drug library using the open-source cheminformatics software RDKit Python API^30^ and performing an all-against-all comparison using the Tanimoto coefficient. Once the features of a molecule have been quantized into a vector, the Tanimoto coefficient is a score of how many bits two vectors have in common divided by the number of bits by which they differ, i.e., |*A* ∩ *B*|*/*|*A* ∪ *B*|, where A and B represent compounds in binary vector form, and |*A*| is the length of the vector.

For efficiency and accuracy, we described our putative drug library using well studied 2D molecular fingerprints.^44^ Specifically, we used Morgan fingerprints,^67^ otherwise known as Extended Connectivity Fingerprints (ECFP, a circular fingerprint), one Functional Class Fingerprint (FCFP, a functional class fingerprint^68^), and fingerprints from RDKit (RDK, a linear fingerprint). Circular fingerprints are bit vector representations of compounds encoding the presence of molecular substructures constructed outward from all starting positions (all atoms) in a radial fashion; functional class fingerprints are binary vectors which encode the presence of predefined “functional” features of a compound; and linear fingerprints encode the presence of molecular substructures built in a linear fashion from all possible staring points (all atoms).^62^

All fingerprints are additionally described by the length of the molecular substructure (“radius” or “diameter” depending on the type and implementation) captured. For instance, ECFP4 is a fingerprint created using ECFP with diameter four. Specific ligand-based pipelines in CANDO are identified according to the molecular fingerprint used, i.e., “ECFP4” refers to the CANDO pipeline where compounds are represented using the ECFP4 molecular fingerprint.

Hert et al. found the optimal results for quantifying relationships between drug classes was achieved using ECFP4 fingerprints with similarities calculated using the Tanimoto coefficient.^59^ We extended this to ligand-based drug repurposing using vectors of 2048 bits instead of the 1024 used in.^59^ We calculated the Tanimoto coefficient between the fingerprints of all possible pairs of the 3733 compounds in our library, and used this to populate a compound-compound similarity matrix, just as we did with the v1 pipeline, allowing us to sort and rank all compounds relative to each other. Fingerprints could not be created for twelve of the 3733 compounds in our putative drug library, which were generally large compounds with metal chelation or long polymers. We then evaluated benchmarking performance of the ligand-based pipelines as described further below.

### Data fusion pipelines

We combined rankings from the v1 pipeline with the new molecular fingerprint rankings using one of the following criteria: lower of two rankings (MIN), higher of two rankings (MAX), sum of two rankings (SUM), average of two rankings (AVG). This is known as “rank-based data fusion”.^69^ We also combined the compound-compound similarity scores from v1 and the ligand-based pipelines using the multiplication of raw similarity scores (MUL), a type of “kernel-based data fusion”. ^69^ After multiplying the similarity scores from two pipelines, the compounds are sorted and ranked based on the newly calculated scores. As in v1 and the ligand-based pipelines, the compound-compound rankings from these data fusion pipelines is then subjected to benchmarking.

### Decision tree pipeline

A goal of CANDO is to make predictions of which compounds are likely to be efficacious against any particular indication. A second is to use analytics to identify causal relationships that predict indication etiology. From the benchmarking, we can determine *a priori* the pipeline that has the best performance for a particular indication, which are then used to generate putative drug candidates for that indication. We constructed a new meta pipeline that makes a decision as to optimal performance on a per indication basis. We made this decision using the top10 average indication accuracy metric (described below), from two choices, v1 and the best performing ligand-based pipeline, namely ECFP4 (see Results). We used this to create a merged set of data which was then benchmarked. For example, the v1 pipeline yields a top10 average indication accuracy of 25% for type 2 diabetes, whereas ECFP4 yields a top10 accuracy of 35%. In the combined decision tree pipeline, we choose to use ECFP4 for the prediction of repurposing candidates for type 2 diabetes, and for the calculation of all benchmarking performance metrics at all cutoffs. We extended this method of choosing the pipeline (between v1 and ECFP4) with higher top10 average indication accuracy to all indications. This aligns with the logic that a clinician or researcher using CANDO can choose the pipeline with the highest accuracy for a particular indication, which is reflected in the benchmarking performance of this combined pipeline.

### Benchmarking pipelines in the CANDO platform

Three measures are used to perform the leave-one-out benchmarking of the CANDO platform pipelines: average indication accuracy, pairwise accuracy, and coverage. Average indication accuracy (%) evaluates the likelihood of capturing at least one drug mapped to the same indication within a particular cutoff from the list of compounds ranked in order of similarity, which is averaged over the 1439 indications with at least two approved drugs and expressed as a percent. Mathematically, this is expressed as *c/d* × 100, where *c* is the number of times at least one other drug approved for the same indication was captured within a cutoff and *d* is the total number of drugs approved for that indication. The top10, top25, top1% (top37), top50, and top100 cutoffs are used, signifying the top ranking 10-100 similar compounds. In other words, the indication accuracy represents the recovery rate of known drugs for a particular indication, which is then averaged across all 1439 indications with at least two approved drugs. Pairwise accuracy (%) is the weighted average of the per indication accuracies based on the number of compounds approved for a given indication. Coverage is the number of indications with non-zero accuracy expressed as a percent.

### Controls

The performance of a given pipeline is evaluated relative to a random control, which is the result that we would expect by chance. The original random control data for v1 was generated by repeated creation of random compound-proteome interaction matrices by sampling from the distribution of values present in the v1 matrix. The benchmarking performance for these random control matrices was calculated as described above and in.^46,63^ However, the new ligand centric pipeline is protein agnostic, and the data fusion ones consist of protein agnostic components. Therefore, we constructed a compound-compound matrix of uniformly random similarity scores to use as controls in this study, i.e., the similarity between any two compounds was assigned a random value between 0 and 1. We sorted and ranked every compound relative to every other compound using this this random compound-compound similarity matrix, and evaluated benchmarking performance as described above.

## Results

### Benchmarking performance of the different pipelines

The new pipelines (Figure 1) generally outperform v1 for all three metrics used to evaluate benchmarking performance: average indication accuracy, pairwise accuracy, and coverage (Figure 2). The MUL:v1,ECFP4 data fusion pipeline, created by multiplying the compound-compound similarity scores (RMSD of interaction signatures) from v1 with the Tanimoto coefficient measured between the compounds described using the ECFP4 molecular fingerprint, yields the overall best performance relative to v1 and the ones based on fingerprint comparisons. Specifically, we obtained the highest top10, top25, and top50 average indication accuracies of 17.3%, 23.8%, and 29.6% using this data fusion pipeline. The highest top1% (or top37) and top100 average indication accuracies of 26.8% and 36.7% were obtained using the pipeline based on the ECFP4 molecular fingerprints. Most of the molecular fingerprint pipelines outperform the original v1 pipeline with the exception of ECFP0, a fingerprint based on simple atom count quantization (Figure 2).

**Figure 2:**
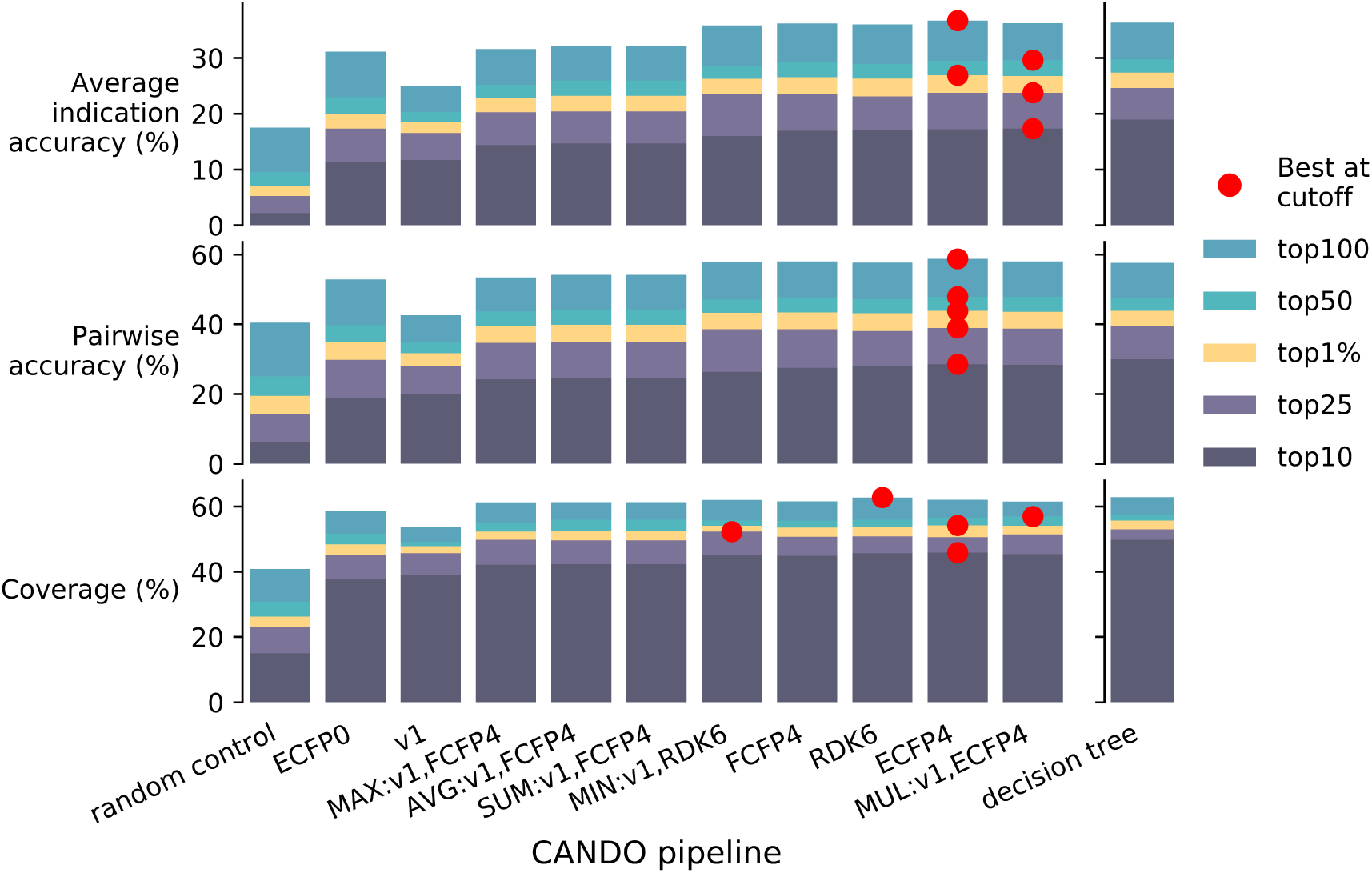
Benchmarking performance of different CANDO platform pipelines. The average indication accuracy (top), pairwise accuracy (middle), and coverage (bottom) for each pipeline are shown at different cutoffs. The value for the top10 cutoff is denoted by dark purple, top25 by light purple, top1% (or top37) by yellow, top50 by green, and top100 by light blue. The individual pipeline with the best performance at each each cutoff is denoted by a red dot. The meta decision tree pipeline was built combining two pipelines, v1 and ECFP4, using the top10 average indication accuracy and so has the highest top10 accuracy and coverage, but is excluded by the “Best at cutoff” marker. The pipelines in all plots are sorted according to increasing top10 average indication accuracy, the most stringent criteria used in our benchmarking. The MUL:v1,ECFP4 pipeline yields the overall best performance relative to the other individual structure-and ligand-based pipelines. The pipeline based on the ECFP4 molecular fingerprint produces the highest top1% and top100 average indication accuracies (top). When assessing pairwise accuracy (middle), ECFP4 is the best performing individual pipeline. The coverage (bottom) plot is a percentage of the 1439 indications for which a pipeline produces a non-zero indication accuracy. The data fusion pipelines of MUL:v1,ECFP4 and MIN:v1,RDK6 have the highest coverage at the top50 and top25 cutoffs, the ECFP4 at the top10 and top50 cutoffs, and RDK6 at the top100 cutoff. Overall, the pipelines using molecular fingerprints have promise and potential for shotgun drug repurposing by themselves, but the data fusion and decision tree pipelines that combine structure-based and ligand-based approaches achieve the best performance while retaining the benefits of both types of approaches.

The decision tree meta pipeline, built by combining other pipelines based on the cor-responding top10 average indication accuracies, yields accuracies of 19.0%, 25.7%, 28.3%, 31.4%, and 39.1% at the five cutoffs used. In contrast, the best performing control generated from uniformly random compound-compound similarity data obtains average indication accuracies of 2.2% at the top10 cutoff, the most stringent one used to benchmark the CANDO platform (Figure 2).

In terms of pairwise accuracy (%), which is the weighted average of the per indication accuracies based on the number of compounds approved for a given indication (see Methods), ECFP4 outperforms all other pipelines, including the decision tree, with accuracies of 28.5%, 38.9%, 43.8%, 47.9%, and 58.8% at the five cutoffs.

The coverage metric evaluates the fraction (or percentage) of the 1439 indications with two approved drugs for which there is at least one instance of a successful recapture or recovery of the known drug within a particular cutoff. The ECFP4 pipeline has the highest top10 and top1% coverage of 45.9% and 54.2%, the MIN:v1,RDK6 yields the highest top25 coverage of 52.3%, the MUL:v1,ECFP4 has the highest coverage at the top50 cutoff of 56.9%, and RDK6 the highest at the top100 cutoff of 62.8%. In contrast, the decision tree pipeline, built in part to increase coverage, has a top10 coverage of 49.8%. This means that for almost half of all the 1439 indications, we capture a drug associated with that indication within the top10 cutoff (Figure 2).

### Distribution of indication accuracies between the two types of pipelines

To compare and contrast the behavior of the structure-based and ligand-based pipelines, we calculated histograms of the average indication accuracies and counts of the highest per indication accuracies at each cutoff for two pipelines (v1 and ECFP4), excluding indications for which a 0% average indication accuracy is obtained. Figure 3 shows that the ECFP4 pipeline has more indications with higher accuracies than v1 (the yellow histogram is shifted to the right of the purple histogram). The Kolmogorov–Smirnov statistical test p-values shown in the corresponding left hand side graph of Figure 3 indicate that the distributions of the v1 and ECFP4 accuracies are drawn from different samples in a statistically significant manner. The Venn diagrams of the 1439 indications in CANDO with more than one approved drug shows that v1 obtains a higher top10 accuracy for 150 indications, while ECFP4 obtains a higher top10 accuracy for 445, and 122 indications have the same non-zero top10 accuracy for both pipelines. As the cutoff increases, more indications have higher accuracies using the ECFP4 pipeline relative to v1, while the number of indications with the same accuracy increases relatively. The orthogonality in the histograms and Venn diagrams indicate that both types of pipelines appear necessary for maximum coverage and accuracy across all the indications. Figure 3 also suggests that additional pipelines and/or improvement in existing pipelines is necessary to recover drugs for ≈500 indications that are not covered by either pipeline at the highest cutoff.

**Figure 3:**
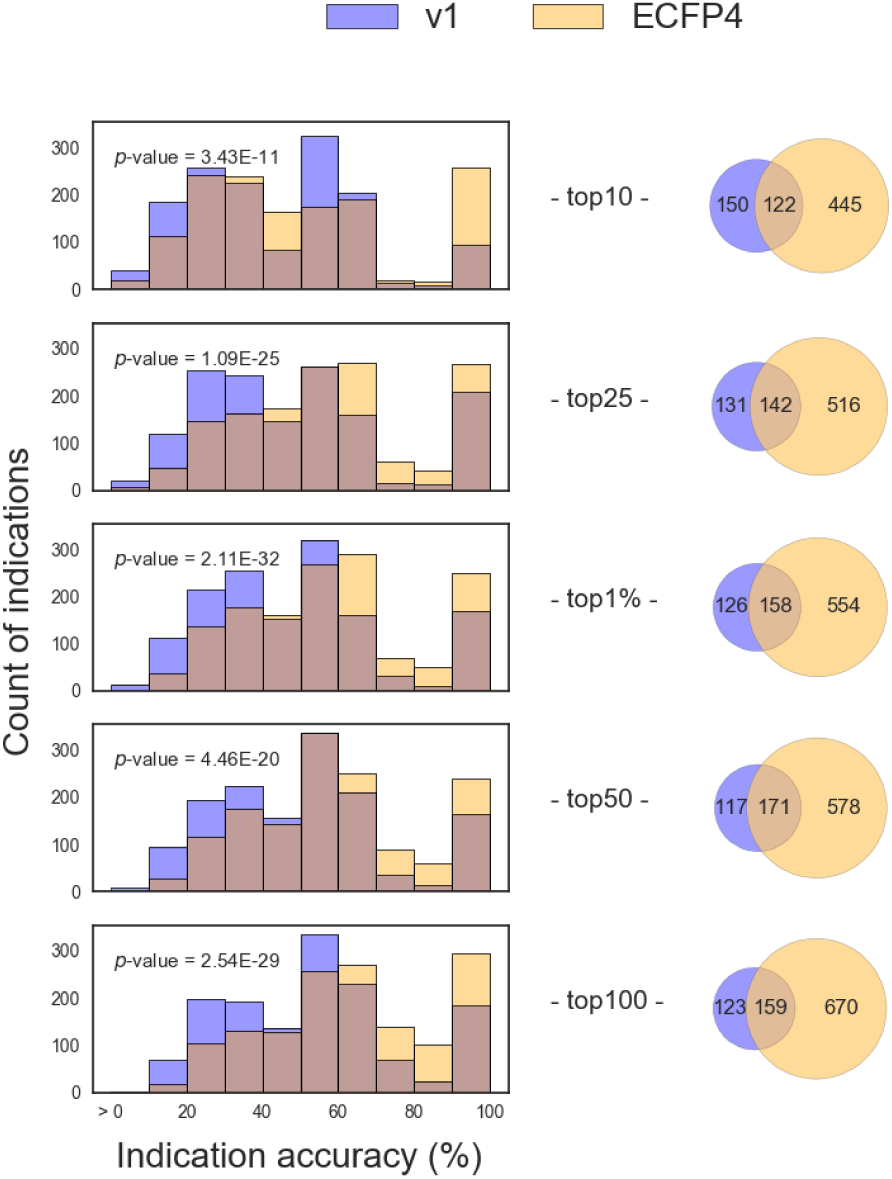
Comparison and overlap of indication accuracy distributions for two CANDO platform pipelines at different cutoffs. The left hand side shows the histograms of the counts of indications with a particular average indication accuracy (or accuracy distributions) for two pipelines, v1 (purple) and ECFP4 (yellow). Indications where both pipelines perform equally well are indicated by brown. For example, at the top10 cutoff, there are approximately 200 indications which achieve an average accuracy between 10 and 20% using the v1 pipeline but just over 100 using ECFP4. At all cutoffs, a greater number of indications with higher accuracies is observed for the ECFP4 pipeline (increase in yellow along the horizontal axis). The *p*-value, derived from the Kolmogorov-Smirnov test statistic applied to the two distributions at each cutoff, indicates that they are significantly different. On the right hand side of the figure are Venn diagrams of the set of indications with higher accuracies at each cutoff (excluding indications with 0% accuracy). For example, at the top10 cutoff, there are 150 indications for which the v1 pipeline yields higher average indication accuracies, 445 for which the ECFP4 pipeline is higher, and 122 with the same performance. The ECFP4 pipeline performs better than v1 for more indications at all cutoffs, but both pipelines appear to be necessary to achieve the best performance across all indications for shotgun drug repurposing.

**Figure 4:**
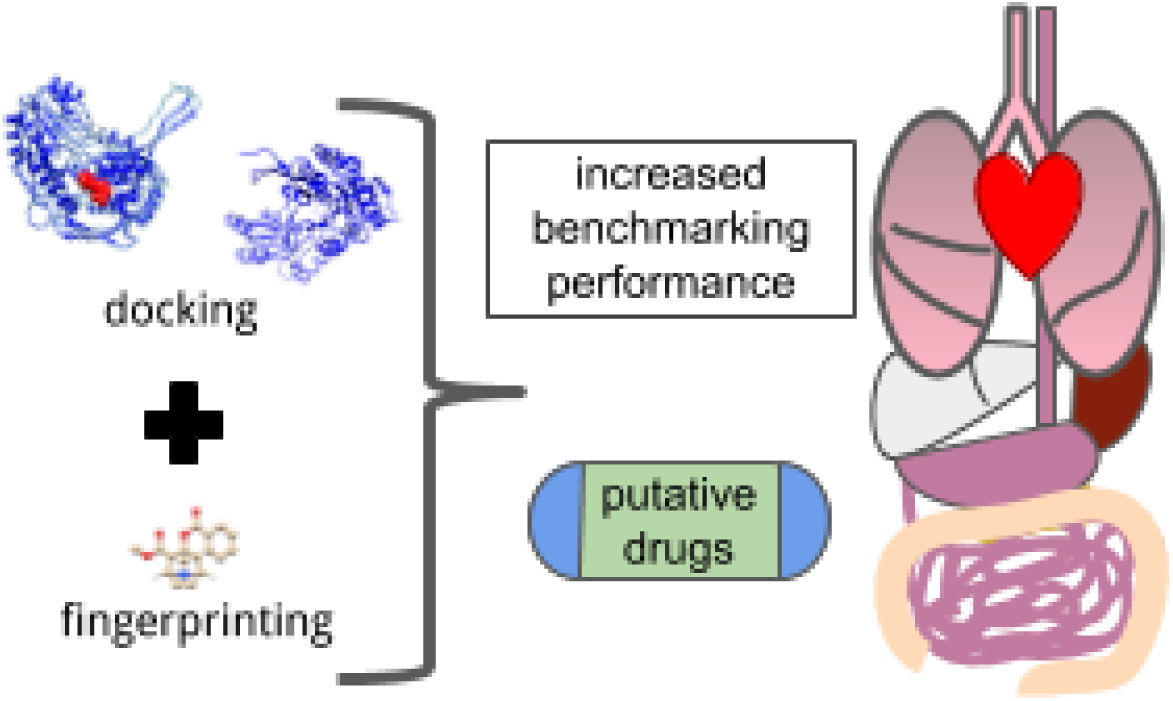
For Table of Contents Only.

### Putative drug candidate generation and validation

The top ranking putative drug candidates generated by the v1 pipeline for eight indications, tuberculosis, malaria, hepatitis B, hepatitis C, systemic lupus erythematosus, type 2 diabetes mellitus, and Alzheimer’s disease, are available from Figure 3 and Supplementary Material of a previous publication.^45^ The top candidates were chosen based on a concurrence score which is “the number of occurrences of particular compounds in each set of top 25 predictions generated for all of the drugs approved for a particular indication”.^45^ Using this concurrence score measure, we generated the top candidate drugs to treat the same indications with the ECFP4 molecular fingerprint and the MUL:v1,ECFP4 data fusion pipelines. We then searched the biomedical literature using PubMed and Google Scholar for published studies corroborating these top candidates.

Both of the new pipelines predict colistin (polymyxin E) as a treatment for tuberculosis, which has been studied as a potentiator of anti-tuberculosis drugs. ^70^ Minocycline was a top result from both pipelines for malaria, which has been shown to protect against certain types and complications.^71^ However, the CDC recommends using doxycycline and not minocycline as malaria prophylaxis.^72^ Additionally, for malaria, both new pipelines list tigecycline among the top ranked candidates, which has shown antimalarial activity in preclinical studies. ^73,74^

All three pipelines recommend known antivirals for hepatitis B. For hepatitis C, all three pipelines list didanosine in the top ranked candidates. Unfortunately, concurrent use of didanosine and traditional hepatitis treatments may induce dangerous consequences for the patient,^75^ illustrating the need for careful expert curation of top candidates generated by the CANDO platform. For Alzheimer’s disease, one of the highest scoring compounds from the MUL:v1,ECFP4 pipeline was dextromethorphan. In 2015, a study was published showing dextromethorphan hydrobromide–quinidine sulfate was well tolerated in patients with Alzheimer’s disease and had clinically relevant efficacy in treating patients, as measured via agitation. ^76^ These examples indicate new putative drug candidate generation by the CANDO platform with these integrated pipelines is likely to work as well, if not better, relative to the prospective validation studies previously done using v1 or its components.^8,50,77–80^ The full list of drug candidates for the above indications based on the concurrence score using the newer pipelines are given in the Supporting Information and available at http://protinfo.org/cando/results/fingerprinting_cando. Putative drug candidate predictions for all 2030 indications in the platform using the v1 pipeline are available at http://protinfo.org/cando/data/raw/matrix/.

## Discussion

### Interpretation of results

Higher benchmarking accuracies are expected to result in better drug repurposing predictions. The top ranked similar compounds to the known drugs for a particular indication using the pipeline with the best benchmarking performance is expected to produce hits and leads with the highest likelihood of success when validated in downstream preclinical and clinical studies. The decreased need to test a large number of compounds with the new pipelines, along with greater confidence in the computational models of drug-indication associations, realizes the goal of drug repurposing: making drug discovery more efficient by reducing the labor, time, and risk in finding new uses for existing therapeutics.

Using the new pipelines based on molecular fingerprinting and data fusion with v1 (Figure 1), we obtain better benchmarking performance than using v1 by itself (Figure 2). Our cutoffs for calculating performance metrics are chosen based on collaborations with wet lab experimentalists willing to test the top candidates generated by our CANDO platform for particular indications. In practice, when working with preclinical and clinical collaborators, we currently employ the decision tree approach of selecting the pipeline with the highest accuracy for a specific indication and the desired cutoff. For example, if a collaborator is capable of validating ten candidates for Precursor B-Cell Lymphoblastic Leukemia-Lymphoma (MeSH identifier D015452), which is one of the 150 where the benchmarking performance is better using the v1 pipeline relative to ECFP4, then we would use the former pipeline to generate the top ten putative drug candidates for this indication.

The new integrated pipelines also yield a higher number of indications covered relative to v1, i.e., more indications with a non-zero accuracy, demonstrating their generalized utility for shotgun drug repurposing. Indication-specific validation studies may rely on the pipeline with highest accuracy for that indication, but CANDO platform development in shotgun drug repurposing requires that the coverage also increase in addition to the average indication and pairwise accuracy. The best performing random control achieves a top10 average indication accuracy of 2.2%, and the random control based on random sampling from the distribution the v1 compound-protein interaction matrix values yielded a top10 accuracy of 0.2%.^45,46^ These random control accuracies are at least an order of magnitude lower than the accuracies obtained using the newer pipelines, and align with expected hit rates in high throughput screening. ^81^ All pipelines yield better performance when compared to the random control (Figure 2), and the differences between the performances of the different pipelines and that of the control signify the value added by our chosen approaches. The orthogonality in the histograms and Venn diagrams of Figure 3 indicate that both types of pipelines appear necessary for maximum coverage and accuracy across all the indications.

### Limitations and future work

We have added new pipelines based on ligand-based fingerprint comparisons to the CANDO platform (Figure 1) that increase benchmarking performance relative to the original v1 protein-centric pipeline (Figure 2). We are further enhancing CANDO by improving the performance of existing pipelines via parameter optimization, ^82^ exploration of different docking approaches to generate the compound-proteome interaction signatures, ^83^ adding new orthogonal pipelines based on compound-pathway signatures, ^63^ implementing more sophisticated data fusion and machine learning approaches, and by continued dissection of the features responsible for pipeline performance and behavior. ^46,47,63^

Notwithstanding the relative benchmarking performance of the existing CANDO platform pipelines, the structure-based virtual screening or protein docking pipelines are not without their merits. The protein-centric approach enables mechanistic understanding of drug action by modeling compound-protein interactions at the atomic level. Additionally, the protein-centric approach readily lends itself to problems in precision medicine/drug repurposing: Incorporating genetic changes, and modeling amino acid mutations due to non-synonymous nucleotide polymorphisms in protein structures, will result in altered compound-protein interaction scores, allowing us to tailor drug repurposing candidates to an individual genome/proteome. The protein-centric approach facilitates consideration of polypharmacy, where the cumulative effects of multiple drugs on protein targets can be evaluated by the analysis and integration of the corresponding drug-proteome interaction signatures, which can then be used to generate putative drug cocktails and combination therapy candidates. The protein-centric pipeline may also be used to generate putative drug candidates for indications without any approved drugs, but where the target protein or proteome is known. ^8^

We are continuing to enhance the virtual screening pipelines to model reality more accurately, with the goal of increasing compound-proteome signature comparison accuracy. For instance, we are exploring the use different molecular docking programs, such as CANDOCK^84,85^ and AutoDock Vina, ^86^ to populate the compound-proteome interaction signatures. An updated version of the v1 pipeline, v1.5, with parameters optimized for scoring compound-proteome interactions, yields benchmarking performance that is 10% higher relatively at the top10 cutoff (12.8% for v1.5 versus 11.7% for v1).^82^ By combining the improved protein centric and protein agnostic pipelines using data fusion, we obtain the best performance and retain the benefits of both types of approaches, while minimizing the weaknesses of any single approach.

The higher benchmarking performance obtained by the ligand-based pipelines may in part be due to the nature of drug discovery and development, which is biased in favor of already effective compounds in an effort to break into a new market or retain market dominance by generating new intellectual property. New drugs are often derivatives of existing ones with small changes. ^87,88^ Repurposing based on molecular fingerprint similarity will be highly enriched for these “me too” compounds, ^88^ given that the approach to shotgun drug repurposing in the CANDO platform is currently based on detecting drug-compound similarities.

Our benchmarking performance metrics are biased toward reporting particular pipelines as better when they capture what is already known/approved, and not novel repurposing candidates which will work to treat or cure an indication in reality. Barring large scale preclinical validation of putative drug candidates, it remains a reproducible and a meaningful measure in our studies.^45–47^

Our goal in this study was to assess the value of adding fingerprinting and data fusion pipelines to the existing protein-centric pipelines in the CANDO platform, and not an exhaustive enumeration, comparison, and fusion of ligand- and structure-based approaches for identifying drug associations. ^89^ More sophisticated fingerprint representations encode the structures of compounds differently and capture unique features particularly of relevance to drug discovery and repurposing. Future work will extend our analyses to include additional fingerprints that can be created using RDKit, including the Long Extended and Feature Connectivity Fingerprints (LECFP and LFCFP, respectively). Longer fingerprints have been shown to better describe a compound with less redundancy, leading to increased accuracy in virtual screening. ^90^

Features and categories of indications, proteins, and compounds all influence the drug repurposing accuracy of CANDO. We are continuing to undertake thorough experiments exploring the roles of particular features responsible for benchmarking performance. ^46,47,63^ Incorporating machine learning to understand how compound-proteome interaction signatures influence performance will help us find the most parsimonious molecular descriptors for compounds. Drugs may have targets beyond proteins, including DNA and RNA.^91,92^ To better model how a compound interacts with all potential targets, we are integrating compound-nucleic acid interaction modeling into CANDO. Finally, we are working with collaborators to validate the predictions from the various pipelines in preclinical and clinical studies, which represents the ultimate test of the CANDO platform.

## Conclusions

CANDO is a computational platform for shotgun drug discovery and repurposing. We implemented new ligand-based and data fusion pipelines in the CANDO platform, and obtained substantial improvement in benchmarking performance using a combination of protein-centric and protein-agnostic methods. These improved results indicate greater confidence in drug repurposing predictions made by us using CANDO and demonstrate the value of considering different, orthogonal, types of approaches for calculating compound-compound similarities. Our integrated approach moves us closer to developing an accurate, robust, and reliable computational drug repurposing platform, and using it to understand how small molecules interact with each other and with larger macromolecules in their corresponding environments.

## Acknowledgement

This work was supported in part by a National Institute of Health Director’s Pioneer Award (DP1OD006779), a National Institute of Health Clinical and Translational Sciences Award (UL1TR001412), and startup funds from the Department of Biomedical Informatics at the University at Buffalo.

The authors thank Dr. Zackary Falls, William Mangione, Matt Hudson, and Dr. Manoj Mammen for their contributions to this project, including software development, and manuscript reading.

## Supporting Information Available

All Supporting Information about this project can be found at http://protinfo.org/cando/results/fingerprinting_cando.

